# Assessing the sensitivity of EEG-based frequency-tagging as a metric for statistical learning

**DOI:** 10.1101/2021.06.01.446686

**Authors:** Danna Pinto, Anat Prior, Elana Zion Golumbic

**Author notes:** <u>Corresponding Author</u>: Elana Zion Golumbic, The Gonda Center for Multidisciplinary Brain Research, Bar Ilan University, Ramat Gan, Israel.

## Abstract

Statistical Learning (SL) is hypothesized to play an important role in language development. However, the behavioral measures typically used to assess SL, particularly at the level of individual participants, are largely indirect and often have low sensitivity. Recently, a neural metric based on frequency-tagging has been proposed as an alternative and more direct measure for studying SL. Here we tested the sensitivity of frequency-tagging measures for studying SL in individual participants in an artificial language paradigm, using non-invasive EEG recordings of neural activity in humans. Importantly, we use carefully constructed controls, in order to address potential acoustic confounds of the frequency-tagging approach. We compared the sensitivity of EEG-based metrics to both explicit and implicit behavioral tests of SL, and the correspondence between these presumed converging operations. Group-level results confirm that frequency-tagging can provide a robust indication of SL for an artificial language, above and beyond potential acoustic confounds. However, this metric had very low sensitivity at the level of individual participants, with significant effects found only in 30% of participants. Conversely, the implicit behavior measures indicated that SL has occurred in 70% of participants, which is more consistent with the proposed ubiquitous nature of SL. Moreover, there was low correspondence between the different measures used to assess SL. Taken together, while some researchers may find the frequency-tagging approach suitable for their needs, our results highlight the methodological challenges of assessing SL at the individual level, and the potential confounds that should be taken into account when interpreting frequency-tagged EEG data.

## Introduction

Statistical Learning (SL) refers to the remarkable ability to implicitly learn the rules and relationship between different stimuli and events in the environment. The capacity for SL has been studied in both humans and non-human species (Santolin and Saffran, 2018; Kang et al., 2021), and has been demonstrated across different sensory domains, emerging relatively early in infancy (Saffran, Jenny R., Aslin, Richard N., Newport, 1996; Saffran et al., 1997; Gómez and Gerken, 2000; Graf et al., 2007; Pelucchi et al., 2009). SL has been hypothesized to play an important role in the development of many key cognitive abilities such as communication skills, object recognition and sensory-motor learning (Read, 1992; Thiessen and Saffran, 2003, 2007; Evans and Robe-torres, 2009; Emberson et al., 2011; Arciuli and von Koss Torkildsen, 2012; Kidd, 2012; Misyak and Christiansen, 2012; Hsu et al., 2014; Spencer et al., 2015; Erickson and Thiessen, 2015; Kidd and Arciuli, 2016; Siegelman et al., 2017). And yet, despite the potentially pivotal role of SL for cognition, current empirical metrics used to assess SL, particularly at the level of individuals, are largely indirect, and often have low sensitivity.

In typical SL experiments a sequence of stimuli is presented in which the transitional probabilities between consecutive stimuli are manipulated so that some items carry predictive information about which stimulus will follow. One prominent example is the Artificial Language Paradigm, where participants hear sequences of syllable-triplets that are always presented consecutively (transitional probability = 1) and thus form “words” in an Artificial Language (which we refer to throughout this paper as *pseudowords*). Participants are exposed to these stimuli for a short period of time (exposure phase), which can range between 2-24 minutes (Saffran et al., 1997; Karuza et al., 2013; Batterink et al., 2015; Franco et al., 2015b), and then perform a test to assess whether the statistical regularities within the sequence have been picked-up by the listener. A variety of explicit and implicit tests can be applied to evaluate SL following an exposure phase, such as 2-alternative forced-choice test (2AFC) or target-detection tasks (Batterink et al., 2015; Batterink, 2017; Batterink and Paller, 2017). Behavioral results on these tests usually show moderate yet above-chance performance when analyzed at the group-level. For example, performance on 2AFC tasks ranges between 54%-68% across studies, which constitutes a significant yet fairly weak demonstration of learning (Saffran et al., 1997; Olson and Chun, 2001; Toro et al., 2005; Turk-Browne et al., 2005; Tyler and Cutler, 2009; Buiatti et al., 2009; Kim et al., 2009; Fernandes et al., 2010; Franco et al., 2011, 2015b; Batterink et al., 2015; Siegelman and Frost, 2015; de Diego-Balaguer et al., 2015; Frost et al., 2015; Batterink and Paller, 2017). However, success rates of individual participants are rarely reported, and the few studies that do include this data find that at least 30% of the participants show no evidence for SL at all and in many individuals behavioral effects are quite small (Cunillera et al., 2008; Romberg and Saffran, 2013; Franco et al., 2015b). It is also worth noting the within-subject correlation between different behavioral tasks (e.g. explicit vs. implicit tests) is often low, raising questions about what is the optimal experimental operationalization for capturing and assessing SL (Batterink et al., 2015, Franco et al., 2015; Misyak et al., 2010b). Given the hypothesized fundamental role of SL for a variety of cognitive processes (Erickson and Thiessen, 2015; Arciuli, 2017) it seems pertinent to develop a more robust empirical measure of SL, that can reliably assess whether or not SL has occurred at the level of individual subjects.

Rather than relying on post-exposure behavioral testing for assessing SL, an alternative approach is to analyze participants’ neural activity during the exposure phase and look for evidence that statistical regularities within the stimulus are picked up. Along these lines, an EEG-based frequency-tagging approach has recently been proposed using a variation of the Artificial Language Paradigm (Buiatti et al., 2009; Batterink and Paller, 2017, 2019; Getz et al., 2018; Batterink, 2020; Choi et al., 2020; Elmer et al., 2021; Henin et al., 2021; Kiai and Melloni, 2021; Lukics and Lukács, 2021). In this version syllables are presented at a constant rate (e.g. *X Hz*), and consequently the tri-syllabic pseudowords also occur at a fixed rate (*X/3 Hz*). These two levels of information are thus distinguishable in frequency, which can potentially be observed in the spectrum of the EEG neural recording. This frequency tagging approach has been successfully employed for studying real speech processing, demonstrating that a peak at the word-level frequency emerges in the spectrum of the neural response when syllables make up words that participants know, but not if they are in a foreign language or do not form recognizable words (Ding et al., 2015; Makov et al., 2017; Har-shai Yahav & Zion Golumbic 2021). Applying this approach to a SL paradigm, Batterink & Paller (2017) demonstrated that the ratio between the power at the syllable vs. pseudoword frequency during the exposure-phase was positively correlated with behavioral performance on an implicit (but not an explicit) behavioral task for assessing SL. This was taken as an indication for the adeptness (and perhaps advantage) of using frequency-tagging to assess SL experimentally, circumventing the need for overt post-exposure behavioral testing. However, despite the promise held by this approach as providing a more direct and objective measure of SL, some of the previous findings raise questions regarding the sensitivity of this measure, particularly at the level of individual subjects. For example, the individual-level data presented by Batterink & Paller (2017) indicate that SL effects were limited only to a subset of participants, with others showing effects in the opposite direction. Moreover, in that study significant effects were also reported when participants listened to random sequences of syllables, where there should not be any SL. As suggested by recent studies, these results may have been somewhat confounded by acoustic contributions to the neural response at the pseudoword-frequency that occur naturally for these type of stimuli (Luo and Ding, 2020; Har-shai Yahav and Zion Golumbic, 2021; van der Wulp, 2021). In particular, a recent re-analysis of the EEG data originally reported by Batterink & Paller (2017), Van dur Wulp (2021) demonstrated that at least some of the reported effects can be explained by variations in place of articulation of different syllables (known as the Obligatory Contour Principle; OCP), rather than by SL of transitional probabilities between syllables. Consequently, without proper controls, the magnitude of the neural response at the pseudoword-frequency might be overinterpreted as only reflecting SL, while the acoustic contribution to this peak is discounted or ignored.

Therefore, it seems that further validation of the frequency-tagging approach is required, and adequate controls implemented, before adopting it as a demonstrably preferable measure of SL. This is an important endeavor not only for furthering our understanding of the potential, and possible limitations, of frequency-tagging for studying SL in humans, but also to assess whether it is an adequate approach for use in clinical conditions (e.g. non-consciousness states etc.) as well as in non-human species, where data analysis typically relies on within-subject effects and not on group-effects. Using a similar experimental design as Batterink et al (2017), the goal of the current study was to test the validity of the EEG frequency-tagging approach for studying SL, while controlling for potential acoustic confounds, as well as its sensitivity for demonstrating SL effects at the level of individual participants.

## Methods

### Participants

Participants were 40 adults (25 female, 35 right-handed), ages 20-38 (mean = 24.78, SD = 3.96). Due to technical issues, EEG data from one participant and behavioral data on the implicit test from 13 participants was lost. All participants reported normal hearing and had no history of psychiatric or neurological disorders and were native Hebrew speakers. They were paid or received course credit for participation. The study was approved by the IRB committee at Bar Ilan University and participants read and signed an informed consent form prior to starting the experiment.

### EEG Recording and Apparatus

EEG was recorded using a 64 Active-Two system (BioSemi) with Ag-AgCl electrodes, placed according to the 10-20 system, at a sampling rate of 1024 Hz. Additional external electrodes were used to record from the mastoids bilaterally and to monitor eye-movements. The experiment was conducted in a dimly lit acoustically and electrically shielded booth. Participants were seated on a comfortable chair and were instructed to keep as still as possible and breathe and blink naturally. Experiments were programmed and presented to participants using PsychoPy (https://www.psychopy.org) (Peirce et al., 2019). Visual instructions were presented on a computer monitor, and auditory stimuli were delivered through in-ear earphones (Etymotic ER-1). Button-press responses were recorded using a serial response-box (Cedrus RB).

### Stimuli

The stimuli consisted of 18 CV syllables recorded in a male voice. Individual syllables were recorded in random order to avoid effects of co-articulation, and only recordings with a flat intonation were used. The recordings were edited offline so that each syllable was precisely 250ms long (silence periods were added if necessary), and their loudness was equated (Audacity software). Additional audio-editing and concatenation of syllables into longer streams was performed in Matlab (Mathworks). The Artificial Language consisted of six tri-syllabic pseudowords *(PaShuDi, SoGuMa, NoMuBe, TuBiPo, GeRoVa, KaLeVi)*, with each syllable appearing in only one pseudoword. Accordingly, the within-word transitional probability was 1 and the between-words transitional probability was 0.2. Given that the modulation spectra of these type of stimuli natural contain acoustic-driven peaks and frequencies besides the syllable rate itself (Luo and Ding, 2020; Har-shai Yahav and Zion Golumbic, 2021; van der Wulp, 2021), we tested the modulation spectrum of several syllable-triplet combinations and selected the combination that yielded the smallest peaks at the pseudoword rate and/or its harmonics as the pseudowords in this experiment (Figure 1). We also confirmed that the pseudowords do not sound similar to known Hebrew or English words.

**Figure 1.**
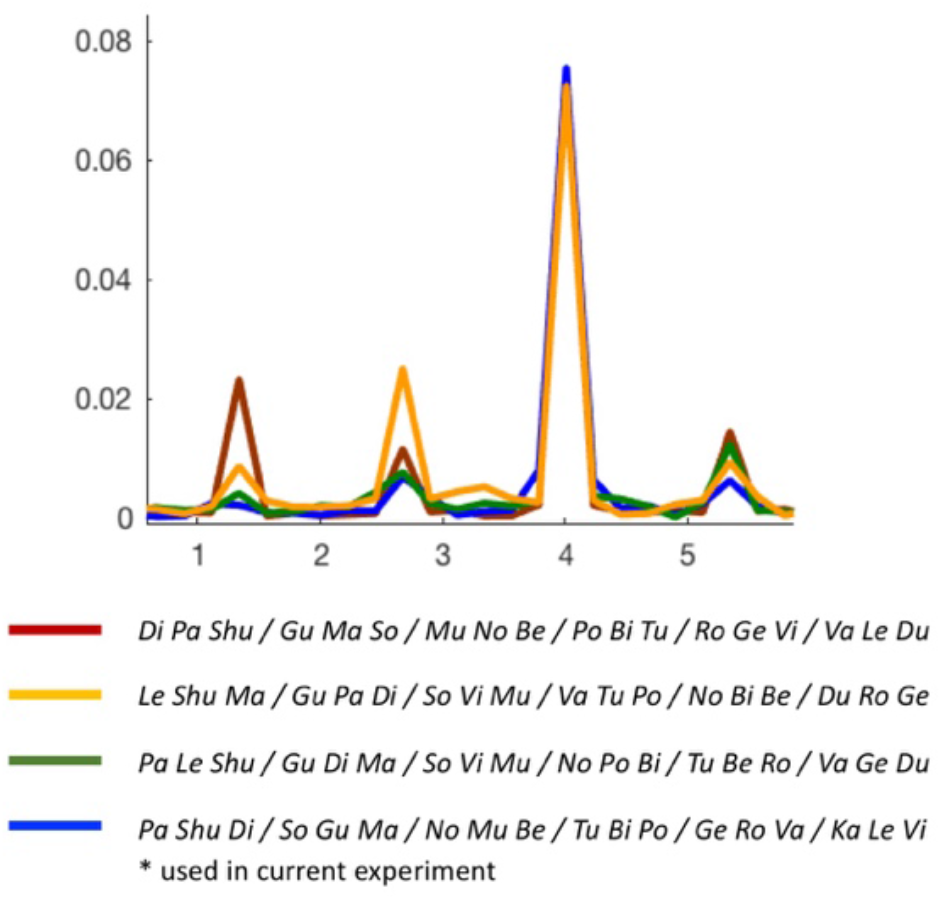
Modulation spectrum for four different versions of the Artificial Language stimuli, all comprised of similar syllables but combined to form different pseudowords. As expected, all stimuli contain a peak at the 4Hz syllable rate. However, as shown here, additional peaks are observed at the pseudoword rate (1.3Hz) and its harmonics, and the magnitude of these peaks varies for the different combinations. As show in similar studies (Luo and Ding, 2020; Har-shai Yahav and Zion Golumbic, 2021, van der Wulp, 2021), these peaks stem from the fact that the same subset of syllables are present in constant positions within the stimulus streams. The Artificial Language stimuli chosen in the current experiment was a combination of syllables that generated relatively small peaks at pseudoword rate frequencies and its harmonics in the modulation spectrum, however these were nonetheless still present (blue line). This motivated the use of Position Controlled stimuli as a means to control for these inherent acoustic peaks, which has a similar modulation spectrum as the Artificial Language stimuli. This allowed us attribute significant differences in the neural response between these two stimuli to effects of Statistical Learning, rather than trivial differences in their acoustic structure.

Since the acoustic-driven contributions to the modulations-spectrum at the triplet-related frequencies could not be fully eliminated from the Artificial Language stimulus, we constructed a position-controlled Baseline stimulus to estimate the extent of these acoustic contributions to the neural signal. The Baseline stimulus consisted of syllable-triplets constructed from the same 18 syllables, but with less consistent transitional-probabilities between them. Similar to the approach used by Makov et al. 2017, in these position-controlled syllable-triplets each syllable maintained the position it held in the original pseudowords, however all possible combinations were allowed (Figure 2, right). This yielded a constant transitional probability of 0.2 both within-triplet and between-triplets in the Baseline stimulus.

**Figure 2:**
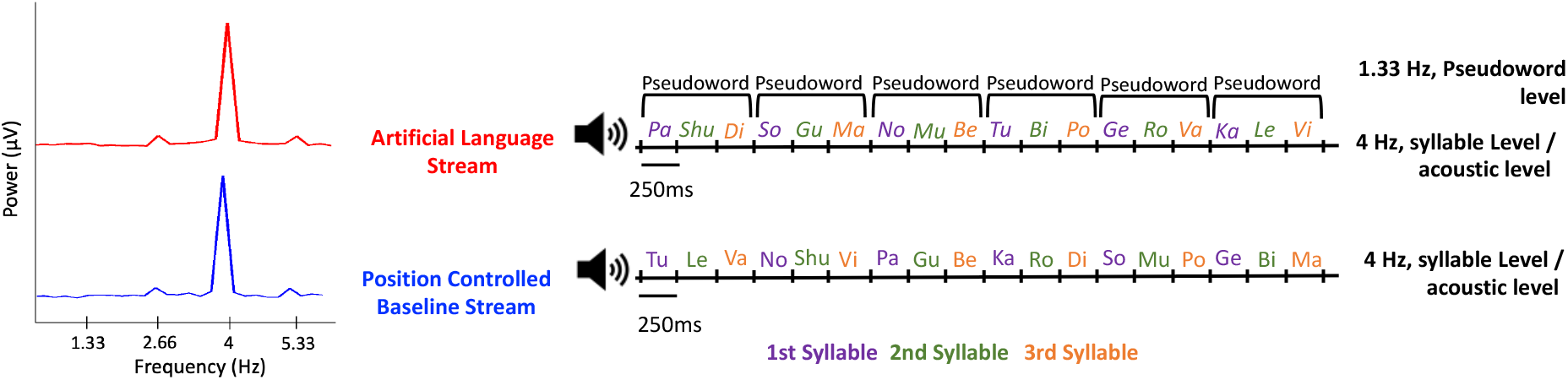
Diagram illustrating structure of the Artificial Language and Position-Controlled Baseline streams used in the current experiment. Left: The modulation-spectrum of the Artificial Language Stream (red) and the Position Controlled Baseline Stream (blue). Both show a prominent peak at the syllable rate (4Hz), as well as more modest peaks at 2.66Hz and 5.33Hz, which are the first and second harmonics of the triplet-rate. Right: Examples of the auditory streams. Both stimuli were comprised of the same syllables, presented at a constant rate of 4Hz. Each stream consisted of syllable-triplets, with each syllable consistently either at the 1^st^ (blue), 2^nd^ (green) or 3^rd^ (orange) position. In the Artificial Language stream, fixed syllable-triplets were used forming pseudoword (within-pseudoword transitional probability = 1; between-pseudoword transitional probability = 0.2), whereas in the Position-Controlled Baseline stream all possible triplet-combinations were used, resulting in a consistent transitional probability of 0.2 between all syllables.

Both the pseudowords and the position-controlled triplets were concatenated to create three 3.22-minute-long streams of the Artificial Language and Baseline conditions. The order of pseudowords and position-controlled triplets in each stream was pseudorandomized, to avoid immediate repetitions of the same triplet and ensure their equal distribution over time. Comparison of the modulation spectra confirmed that this approach resulted in similar peaks at 1.33Hz for both the Artificial Language and Baseline streams, making them highly comparable acoustically and allowing us to gauge the effect of within-word transitional probabilities on the 1.33Hz peak in the neural response, above and beyond any potential acoustic contributions from the stimulus itself (Figure 2, left). Both the Baseline and Artificial Language streams included a 5 second ramping up/down period, to avoid inadvertent cues about syllable positions or pseudoword boundaries.

### Experimental Procedure

#### Exposure Phase

The experiment consisted of several stages. It started with a Baseline exposure stage during which participants listened to the Baseline condition streams of concatenated syllables described above. These consisted of hearing three 3.22-minutes long streams (separate blocks, with breaks between them; total exposure time: ∼10 minutes). Participants were instructed to listen passively to the stimuli with their eyes open and fixated on a point on the screen. In this stage no additional instructions were given. After a brief break during which they completed an English vocabulary test, they were exposed to the Artificial Language streams. Here participants were explicitly told that the stream is made up of words in an Artificial Language, which they are requested to learn for subsequent testing. However, participants were not told the length or number of the pseudowords. The order between exposure-phases was held constant to avoid carry-over learning effects in the Baseline condition after exposure to the Artificial Language.

#### Testing Phase

The Testing Phase consisted of two behavioral tests:

##### 2-Alternative Forced Choice task (2AFC)

The explicit 2AFC discrimination task was designed to follow the commonly used procedure for explicit testing of Statistical Learning (Saffran et al., 1997; Toro et al., 2005, 2011; Buiatti et al., 2009; Tyler and Cutler, 2009; Fernandes et al., 2010; Franco et al., 2011, 2015b; Wang and Saffran, 2014; Batterink et al., 2015; Batterink and Paller, 2017). In addition to the six pseudowords that made up the Artificial Language, six additional part-words were created consisting either of the last two syllables of one pseudoword combined with the first syllable of another, or the last syllable of one pseudoword combined with the first two syllables of another. As such, these are combinations that participants could have heard occasionally during the learning phase, but not as frequently as the actual pseudowords. In each trial, one pseudoword and one part-word were played (random order), and participants were required to indicate via button-press which one was familiar to them from the Artificial Language learning phase. This test consisted of a total of 36 trials (all possible pairs of pseudowords and part-words).

Group-level statistical analysis of performance on the 2AFC task consisted of a single sample t-test testing whether accuracy rates were significantly higher than chance (50%), as commonly done in similar studies (Saffran et al., 1997; Toro et al., 2005, 2011; Buiatti et al., 2009; Tyler and Cutler, 2009; Fernandes et al., 2010; Franco et al., 2011, 2015b; Wang and Saffran, 2014; Batterink et al., 2015; Batterink and Paller, 2017). However, since the 2AFC task consists of only 36 trials and does not necessarily meet the assumptions required for a t-test, we further simulated the null distribution of our specific experiment using a permutation test. We simulated a random 2AFC guessing pattern for 36 trials and calculated the “random hit-rate” of that simulation. This procedure was repeated 1000 times, producing a null-distribution reflecting the probability of achieving a particular hit-rate “by chance” (Figure 3A, shown in gray). Furthermore, we assessed the significance of performance in individual participants by comparing their accuracy rates to a binomial distribution for 36 2AFC trials (Franco et al., 2015b; Siegelman et al., 2017), allowing us to establish which participants showed evidence for Statistical Learning according to the 2AFC test.

**Figure 3.**
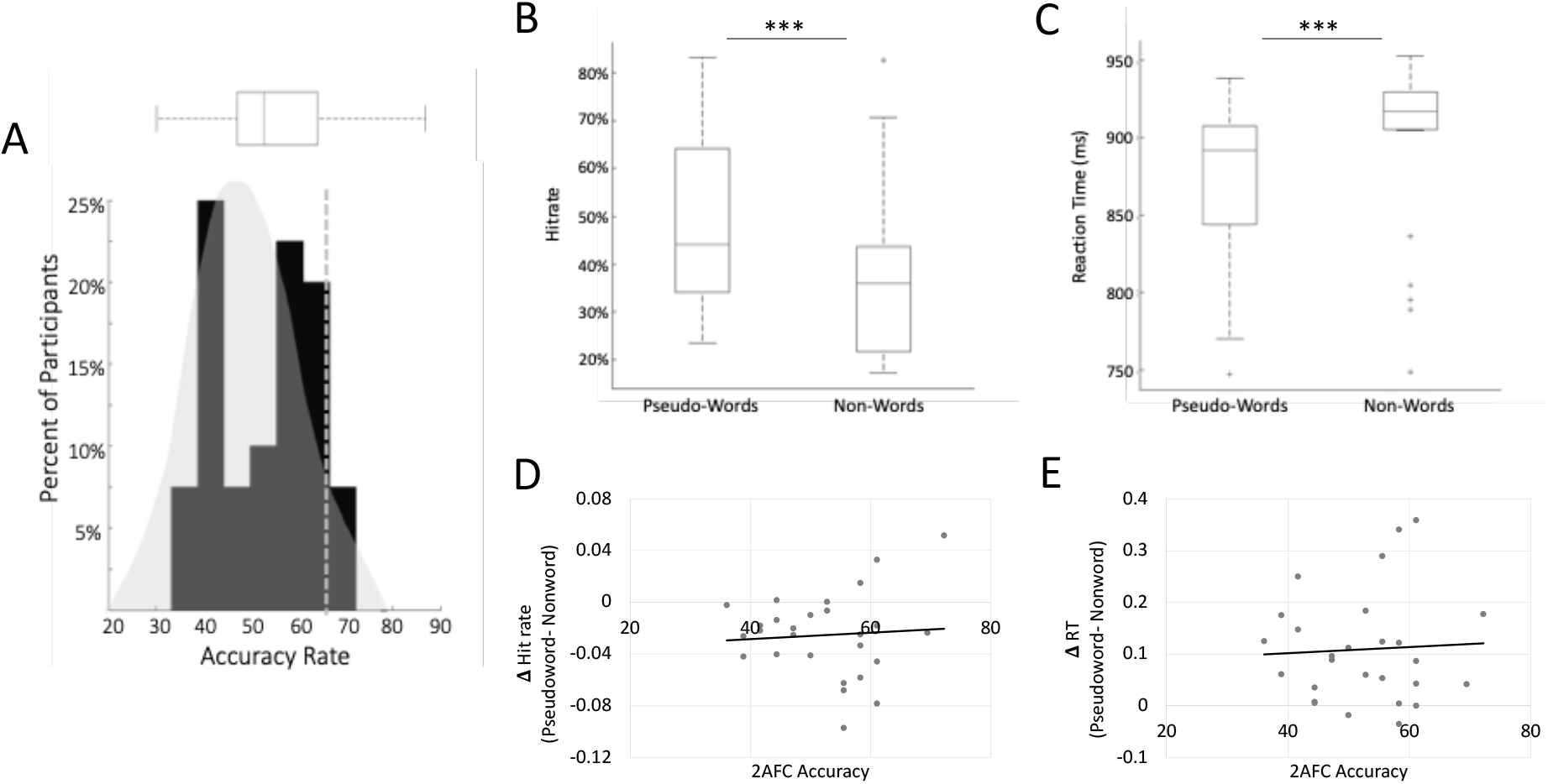
Behavioral Results. A) 2AFC results: Histogram of accuracy rates on the 2AFC task across all participants(black), overlaid on the background of the a-priori binomial distribution of chance-level results in the current design (gray). Top: Interquartile range and group-median of 2AFC results. Dashed gray line: the p=0.05 cutoff for determining whether individual level performance was significantly above chance (relative to the null-distribution). B&C) Target-detection results: Inter-quartile range and group-level median for hit-rates and RTs in response to target-syllables that occurred in the 3^rd^ position of pseudowords vs. target that occurred in Non-words. For both metrics, performance was improved for targets within pseudowords, as indicated by the asterisks (p<.001 between the conditions). Outliers are indicated by a gray plus(+) symbol. D&E) Scatter plots depicting the within-subject relationship between performance on the 2AFC task and the Target-Detection task. No significant correlation was found between accuracy on the 2AFC task and the magnitude of the behavioral effects in the Target-Detection task (difference in hit-rate / RTs for targets in pseudowords vs. targets in non-words).

##### Target detection task

The explicit test was followed by an implicit Target-Detection task, designed based on previous studies using a similar approach (Batterink et al., 2015; Batterink and Paller, 2017). In each trial one syllable was designated as the target and was played twice for participants to familiarize themselves with the sound (e.g. “**Va”**). Then a sequence of syllables was played, and participants were required to press a button when they heard the target syllable. The sequences contained pseudowords from the Artificial Language as well as other triplet syllable combinations (‘non-words’). The target syllable in each trial (e.g. “**Va”**) was placed strategically within the sequences and could occur either as the 3^rd^ syllable of a pseudoword presented in the exposure phase (e.g. GeRo**Va**) or as the 1^st^ or 3^rd^ syllable in a non-word (e.g. **Va**ShuPo or PaMu**Va**). In this task, enhanced target-detection performance for targets presented as the 3^rd^ syllable of a previously learned pseudoword (vs. syllables in a non-word) would serve as an indication that participants had successfully learned the structure of the Artificial Language because they are able to anticipate the target syllable.

For this task, syllables were presented at a constant rate of 2Hz with each trial lasting 22.5 sec and including 4-8 targets. The entire task consisted of 24 trials (4 trials per target syllable). A button press was considered a ***hit*** if it fell within 1 second after the presentation of a target syllable. Otherwise, it was considered a ***false alarm***. Group-level statistical analysis consisted of paired t-tests of hit-rates and Reaction times (RTs) for targets occurring within pseudowords vs. non-words (responses to targets occurring in 1^st^ and 3^rd^ position of non-words were grouped together, since we found no differences between them).

Statistical analysis at the level of individual participants was conducted using permutation tests. The permutation test consisted of random re-labeling of all the responses of a particular participant into two random-conditions, regardless of their original status as pseudowords/non-word targets and taking the difference between the means of the two random-conditions. This procedure was repeated 1,000 times and the differences of the means extracted from each permutation were used to form a null-distribution for each participant. We then took the real difference between the pseudoword and non-word targets in the original data and compared it to the null-distribution. The difference between conditions was considered significant if the real value fell in the top 5%^tile^ of the null distribution (one-tailed). This procedure was performed for both accuracy and reaction times (RT) data.

We further tested whether performance on the two behavioral tasks was correlated, by calculating the Pearson correlations between explicit 2AFC accuracy rates and the implicit Target-Detection task (differences in hit-rates / RTs for targets occurring within pseudowords vs. non-words).

### EEG Data analysis

#### Preprocessing and Spectral Analysis

All EEG preprocessing and analysis was performed in Matlab (The Mathworks) using the toolbox FieldTrip (Oostenveld et al., 2011) as well as custom written scripts. Raw data was first visually inspected for gross movement artifacts which were removed. Eye movements and blinks were corrected using independent component analysis (ICA), and any remaining artifacts were further removed. Noisy electrodes were identified visually and replaced with the weighted average of their neighbors using an interpolation procedure.

The clean data was analyzed separately for the Baseline and Artificial Language exposure blocks. The continuous data was segmented into 4.5 second epochs, which correspond to 6 syllable-triplets. Critically, these segments were perfectly aligned such that they all started with the onset of a triplet. Inter-trial Phase Coherence (ITPC) was used to analyze the neural response at specific frequencies. ITPC was calculated as follows: The Fast Fourier Transform (FFT) was calculated for each individual segment between 0.3-6 Hz using a Hanning window. The phase component at each frequency was used to calculate the ITPC, which is the sum (absolute value) of the phases across segments, as follows:

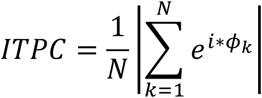

ITPC analysis was performed separately for the Baseline and Artificial Language exposure blocks and was calculated across all blocks as well as separately for each of the three exposure blocks per condition.

Statistical analysis of EEG data focused a-priori on four frequencies of interest: the syllable-presentation rate: 4 Hz; the pseudoword rate: 1.33Hz; and its two harmonics: 2.66 Hz and 5.33 Hz. We tested for differences between the Baseline and Artificial Language conditions at each of these frequencies, both at the group level as well as at the level of individual subjects.

#### Group-level analysis

To avoid a-priori assumptions as to which electrodes would show an effect of Statistical Learning, we performed statistical tests both on the average ITPC across all electrodes (one-way paired t-test, and dependent-sample Bayesian analysis; JASP Team, 2019), as well as at each electrode individually (one-way paired t-test with FDR correction for multiple comparisons). Finding significant peaks at 1.33Hz and/or its two harmonics 2.66Hz and 5.33Hz in the Artificial Language condition would serve as an indication that the statistical regularities within the Artificial Language had indeed been identified and the stream was parsed correctly into tri-syllabic pseudowords. In these analyses, the direction of the expected results was determined a-priori based on the results of previous findings (Buiatti et al., 2009; Batterink and Paller, 2017, 2019), predicting increased responses in the Artificial Language vs. Baseline condition at frequencies associated with pseudoword rate and its harmonics (1.33 Hz, 2.66 Hz and 5.33 Hz), and the reverse pattern at the syllable rate (4Hz).

We additionally tested whether ITPC changed over the course of the exposure stage, using a linear regression analysis. For this we calculated the average ITPC across all electrodes separately for each of the three blocks in each exposure condition, and at each frequency of interest. Average ITPC values for each participant per block were fit using a linear regression model, as implemented in R’s glmer function (lme4 package) (Bates et al., 2015), with Condition (Artificial Language vs. Baseline) and Block (1-3) as fixed effects and Participant as a random effect (model: ITPC = Condition * Block + (1|Participant)). Evidence for SL should manifest as a significant interaction between Condition and Block indicating that ITPC increases systematically across blocks in the Artificial Language condition, but not in the Baseline condition. We used Helmert Forward contrast coding which compares the level of each variable with the mean of all levels of that variable. The regression model was applied separately to the data at each frequency interest.

#### Individual-level analysis

One of our primary goals was to assess whether evidence for Statistical Learning can be gleaned at the level of individual participants using this frequency tagging method. To this end, we used within-subject permutations to test for significant differences in ITPC between the Baseline and Artificial Language conditions. For each participant, we randomly switched the label of half the epochs between the Baseline and Artificial Language conditions and calculated the ITPC-difference between these two null-conditions at each of the four frequencies of interest (averaged across all electrodes). This was repeated 1000 times to create a null-distribution, and if the real difference in ITPC between conditions fell within the top 5% of this null-distribution it was considered significant (p=0.05, one way). This procedure was performed for each participant at each frequency of interest. Lastly, we also tested for correspondence between the EEG-based metric and the behavioral metrics.

## Results

### Behavioral Results

Accuracy levels on the 2AFC task at the group level were not significantly different than chance [mean=52.56%, SD=0.10, t(38)= 1.60, p=.12]. Moreover, when comparing accuracy rates at the individual-level to the a-prior binomial distribution of accuracy rates expected by chance (Figure 3A we find that only N=3 participants (7.5%) had performance that fell in the top 5^th^ percentile of the chance-level distribution (cutoff = 65%, p=0.05).

Due to technical failures, data from the Target Detection task was available only for 27 out of the 40 participants. In the remaining subset, we found significantly higher accuracy and faster RTs for target syllables the occurred in the 3^rd^ position of a pseudoword vs. targets that were part of a non-word [hit-rates: t(26)= 5.35, p<.001; RTs: t(26)=-4.05, p<.001] (Figure 3B&C). Statistical analysis of behavioral results at the Individual-level analysis found that N=19 participants (70%) had a significant effect on *either* hit-rate or RT, however only N=3 participant showed significant effects for *both* measures.

When comparing individual-level results on the two tasks within participants, we failed to find any significant correlations between accuracy rates on the 2AFC task and the magnitude of the behavioral effects in the Target-Detection task (difference in hit-rate / RTs for targets in pseudowords vs. targets in non-words; hit-rate: Pearson’s r^2^=0.05, p=0.79; RTs: r^2^=0.07, p=0.72; Figure 3D&E).

### EEG Results

#### Group-level Analysis

Figure 4A shows the mean ITPC spectra across all participants and electrodes, across the two conditions. Consistent with previous effects of SL, ITPC at the pseudoword rate (1.33Hz) was significantly larger in the Artificial Language condition vs. Baseline [t(38)=2.01, p=0.023; BF=2.057 (anecdotal support)], and an even stronger enhancement was found at the 3^rd^ pseudoword rate harmonic [5.33Hz; t(38)=2.881, p=0.003; BF=11.893 (strongly supported)], although the peak at the 2^nd^ pseudoword rate harmonic (2.66Hz) was not significantly different between conditions [t(38)=0.804, p=0.213; BF=0.36 (moderate acceptance of H0)].

**Figure 4:**
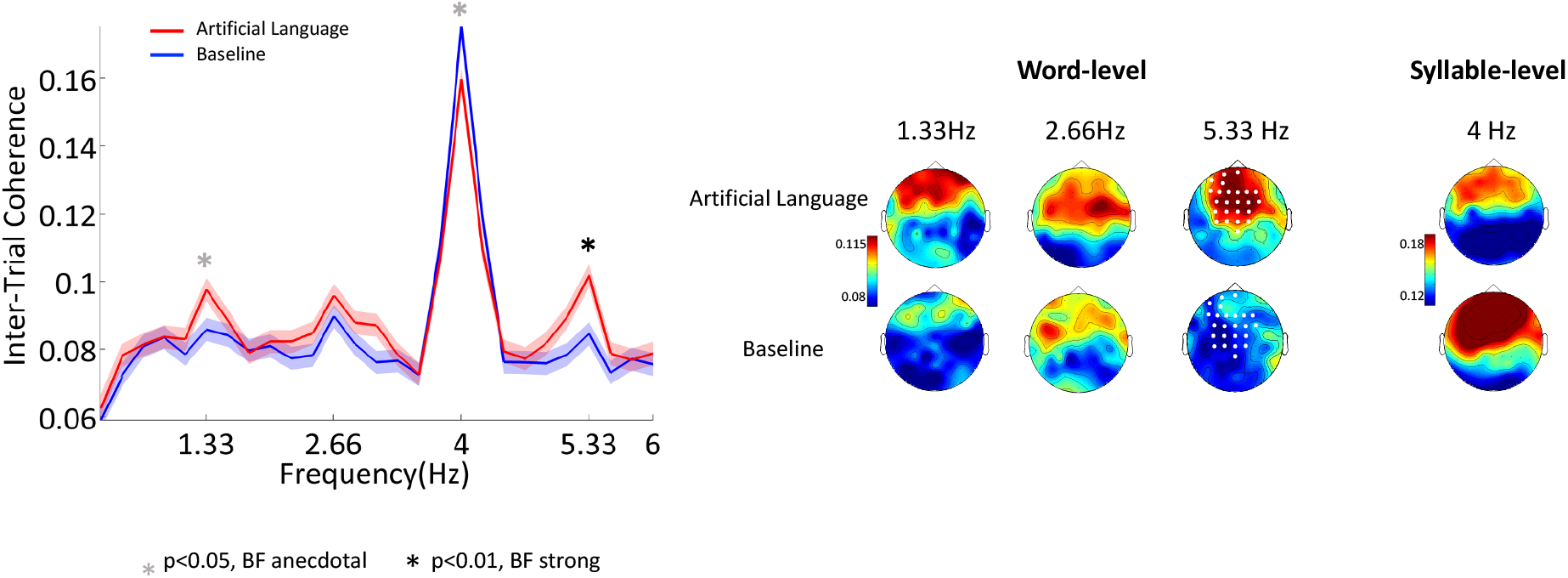
ITPC results. A) Grand average of the ITPC spectrum in the Artificial Language (red) and Baseline (blue) conditions, averaged across all electrodes. Shading indicates SEM across participants (n=39). Asterisks indicate peaks where there was a significant difference between conditions at the group level (t-test p<0.05), accompanied by either strong (black) or anecdotal (gray) BFs. B) Scalp topographies of the ITPC response at each of the four frequencies of interest. White dots indicate electrodes where a significant difference was found between the Artificial Language and Baseline conditions, when comparisons were performed separately at each electrode (fdr corrected).

The ITCP at the Syllable rate (4Hz) was also modulated by the stimulus type but in the opposite direction, with a reduced peak in Artificial Language condition vs. Baseline [t(38)= -1.858, p=.035; BF=2.057 (anecdotal support)].

When repeating the statistical analysis at each electrode separately, the only frequency where a significant effect was found between conditions (after FDR correction) was the 3^rd^ pseudoword rate harmonic (5.33Hz). The effect at this frequency was observed in 24 mid-central electrodes, indicated in the topographic map in Figure 4.

#### By-Block linear regression analysis

Besides looking at the ITPC across the entire experiment, we also performed a linear-regression analysis to test for changes in the response across the three exposure blocks (Figure 5). The main effects of condition (Artificial Language vs. Baseline) reported above for responses as 1.33Hz and 5.33 Hz, were confirmed in this analysis as well [F(190)=2.25, p=0.03, and F(228)=2.12,p=.03 respectively]. However, the effect of block and interactions between condition and block were not significant at any of the frequencies of interest (see Table 1 for full statistical results). Rather, all the effects of condition seem to be present already in the first exposure block and were not further enhanced over time.

**Table 1:** Summary of Statistical Results of Block Analysis. Significant results are indicated in bold and an asterisk.

| Contrast | Frequency | $\beta$ | t | p |
| --- | --- | --- | --- | --- |
| Block Number (*1 vs. 2*) | Word | -0.007 | -0.88 | 0.38 |
|  | Syllable | 0.02 | 1.98 | 0.05 |
|  | 2<sup>nd</sup> Harmony | -0.004 | -0.49 | 0.63 |
|  | 3<sup>rd</sup> Harmony | -0.006 | -0.78 | 0.43 |
| Block Number (*1&2 vs. 3*) | Word | .003 | 0.34 | 0.74 |
|  | Syllable | 0.006 | 0.48 | 0.63 |
|  | 2<sup>nd</sup> Harmony | 0.01 | 1.06 | 0.29 |
|  | 3<sup>rd</sup> Harmony | 0.005 | 0.51 | 0.61 |
| Condition (*Artificial Language vs Baseline*) | Word | 0.11 | 2.25 | **0.03\*** |
|  | Syllable | -0.02 | -2.15 | **0.03\*** |
|  | 2<sup>nd</sup> Harmony | 0.002 | 0.40 | 0.69 |
|  | 3<sup>rd</sup> Harmony | 0.01 | 2.10 | **0.04\*** |
| Interaction<br>*Block Number (1 vs. 2) x Condition* | Word | 0.01 | 1.163 | 0.25 |
|  | Syllable | -0.006 | -0.40 | 0.69 |
|  | 2<sup>nd</sup> Harmony | 0.008 | -0.64 | 0.52 |
|  | 3<sup>rd</sup> Harmony | 0.01 | 1.12 | 0.26 |
| Interaction<br>*Block Number (1&2 vs 3) x Condition* | Word | 0.01 | 1.07 | 0.28 |
|  | Syllable | 0.01 | 0.76 | 0.45 |
|  | 2<sup>nd</sup> Harmony | -0.005 | -0.32 | 0.75 |
|  | 3<sup>rd</sup> Harmony | -0.01 | -0.88 | 0.38 |

**Figure 5:**
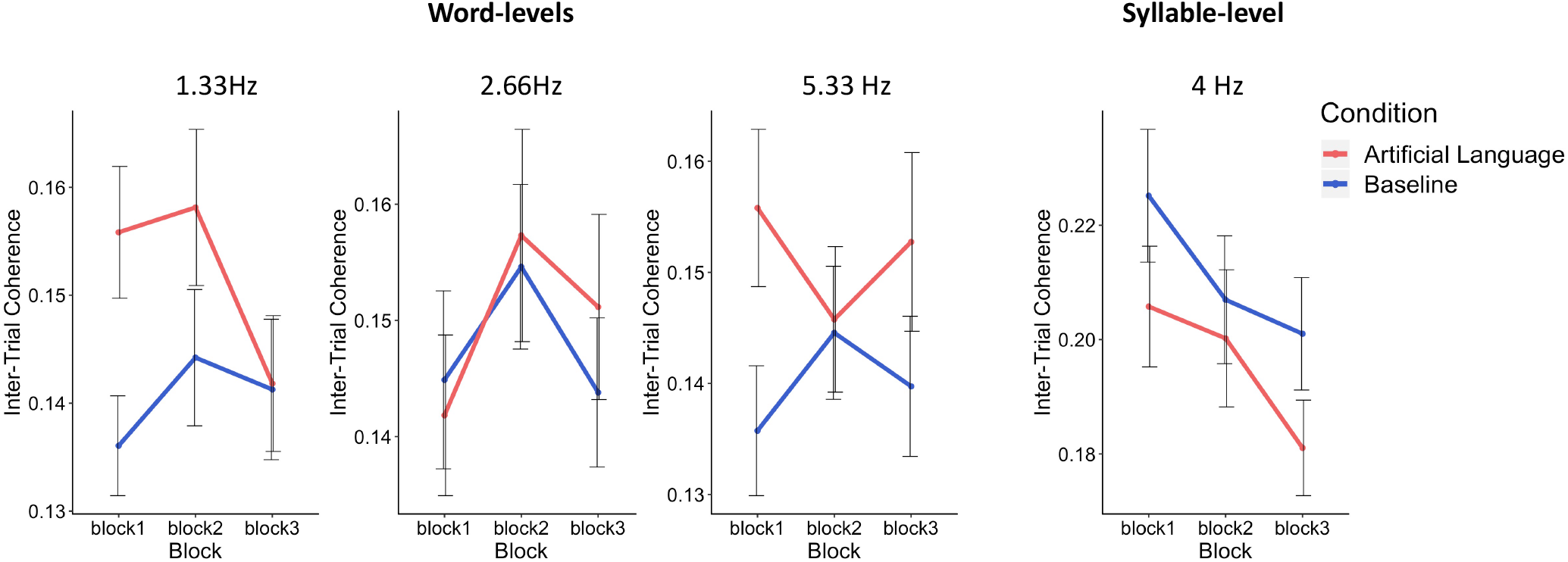
Results of By-Block Analysis: Mean ITPC across the three exposure blocks of each condition, averaged across all electrodes, at each of the frequencies of interest. Linear regression analysis confirmed the main effect of condition at 1.33Hz and 5.33Hz (Artificial Language condition > Baseline), but neither the effects of block nor the interaction between condition and block were significant, at any of the frequencies.

#### Individual Participant Analysis

Assessment of SL effects from the neural response at the level of individual participants, was conducted using permutation tests. Since SL effects could potentially manifest either at the pseudoword rate itself or at any of its harmonics, this analysis was performed at all the frequencies of interest. Overall, we found significant effects of condition in 12/40 participants (30%), with larger responses in the Artificial Language condition vs. Baseline. Of them, in n=5 the effects were at 1.33Hz, in n=3 participants the effects were at 2.66Hz and n=4 participants had an effect at 5.33Hz. Only one participant had significant effects at more than one frequency. In addition, the reduced response in the Artificial Language condition at the Syllable rate (4 Hz) that was observed at the group level, was found to be significant n=6 participants. For examples of the ITPC spectrum of individual participants see Figure 6).

**Figure 6:**
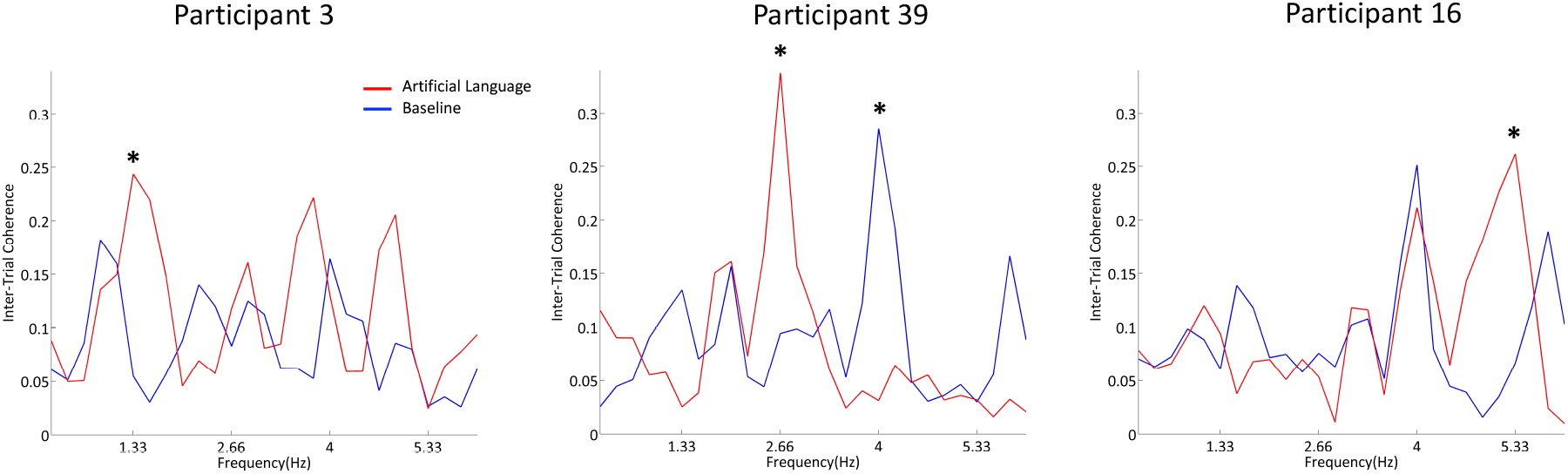
Examples of ITPC spectral of individual Participants: ITPC spectra in the Artificial Language (red) and Baseline (blue) conditions from three participants who showed significant differences between the Artificial Language and Baseline conditions, albeit at different pseudoword related frequencies (indicated by an asterisk). Spectra from each participant are shown from the electrode showing the largest significant effect.

#### Correspondence between ITPC effects and behavior

We next tested whether the ITPC response to pseudoword rates in the Artificial Language condition corresponds to performance on behavioral tasks administered post-exposure. Since the Individual-level analysis revealed inconsistencies in the specific pseudoword related frequencies where significant differences were found across participants (i.e., at the pseudoword frequency itself or one of the harmonics), this prevented us from performing a simple correlation analysis between the ITPC at a particular frequency and behavioral measures. To overcome this between-participant variability, we used two different approaches.

First, we took the average ITPC in the Artificial Language condition across the three pseudoword related frequencies (1.33Hz, 2.66Hz and 5.33Hz), and calculated the Pearson correlations with each behavioral measure. This did not, however yield any significant results (correlation with 2AFC accuracy: r^2^=0.16, p=0.33; correlation with target detection hit-rate: r^2^=0.13, p=0.50), correlation with target detection RTs: r^2^=-0.05, p=0.81).

Second, we separated the participants into two groups based on whether there was evidence for SL from their neural data (regardless of the frequency where this effect was observed) and compared the behavioral results between the two groups. We used the Welch’s Test for Unequal Variance, to account for the different sample sizes in the two groups. In this analysis we found that the group in which significant pseudoword EEG responses were observed also had significantly larger behavioral effects in the Target-Detection task (Figure 7, bottom panel). Specifically, this group had larger differences in RTs between targets occurring in the 3^rd^ position of pseudowords vs. targets within non-words was larger [t(14)= 2.15, p=0.03; BF10=4.18 (moderate support)]. However, this effect was not significant for hit-rates in the Target-Detection task [t(10)=-1.02,p=0.17] or for performance on the 2AFC task [t(21)=-0.19, p=0.85] (Figure 7, top and middle panels).

**Figure 7:**
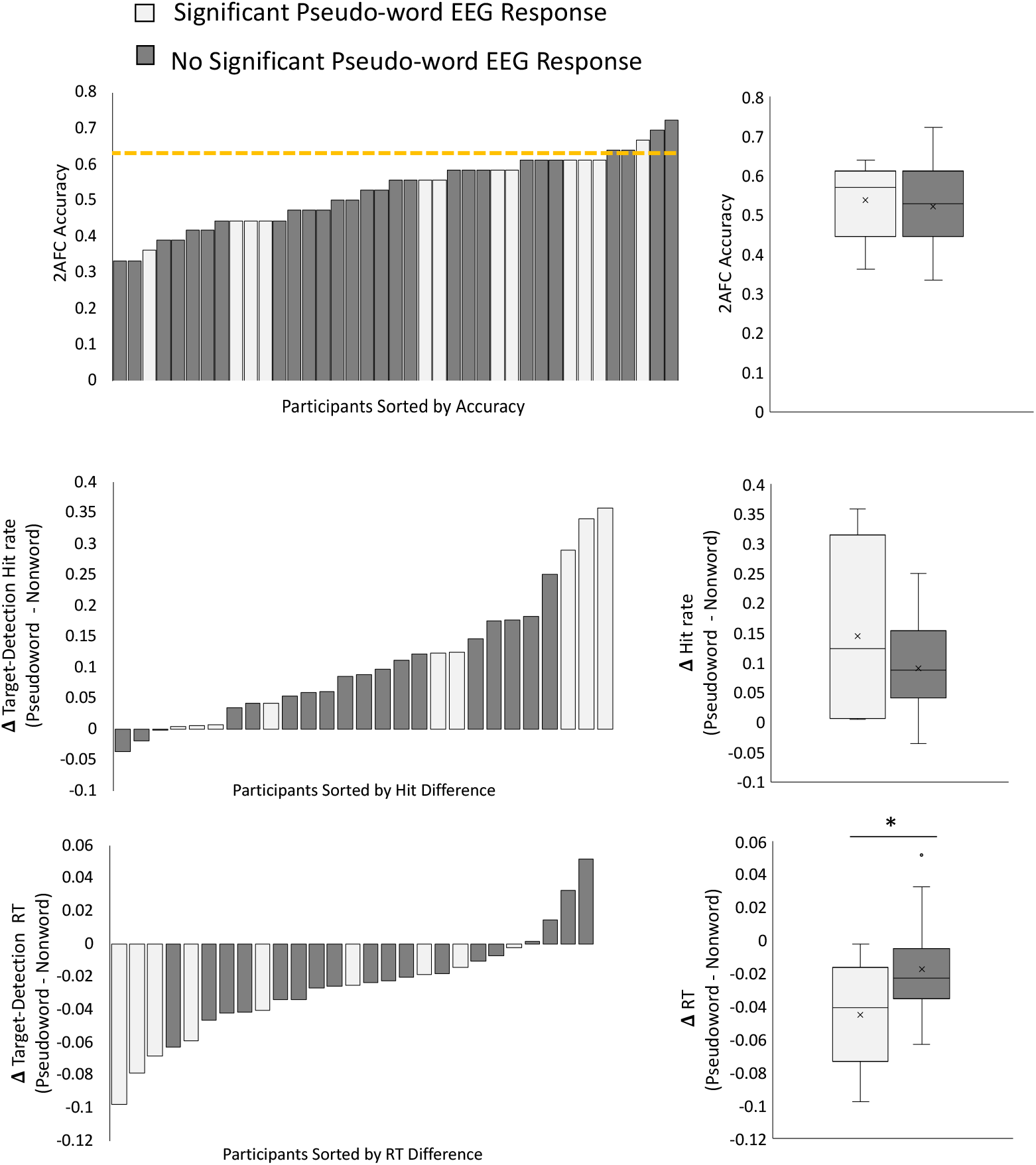
Correspondence between and pseudoword EEG response during the exposure period and post-exposure behavioral tasks. Participants were divided into two groups based on whether a significant response to pseudowords was found in their neural response during the exposure period (gray) or not (black). The left-hand panels show the results of all participants on each of the three behavioral measures: 2AFC accuracy, effects size on hit-rate and RTs in the Target-Detection task (sorted by effect size and color-coded by group). The right-hand panels show the means and SEM of these behavioral measures, in each group. A significant difference between-groups was found for Target-Detection RT effect (bottom panel, p<0.05), but not for the other behavioral measures.

## Discussion

In this study, we tested the sensitivity of the EEG frequency-tagging approach as an online measure for assessing auditory Statistical Learning of an Artificial Language, at both the group-level and within individual-participants. We find that, even after controlling for potential acoustic contributions to the pseudoword frequency, there is still a significant difference between the Artificial Language and the Baseline conditions at the group level. This effect manifested most robustly at the 3^rd^ pseudoword-level harmonic (5.33Hz), and less reliably at the pseudoword-level rate itself (1.33Hz). The previously reported decrease at the 4Hz syllable-level for Artificial Language stimuli was also observed here, but again with low statistical reliability. Effects were observed already during the first exposure block, (3.22 minutes) and did not change significantly with additional exposure. These results help validate the use of the frequency-tagging approach for assessing SL <u>at the group level</u>, while highlighting important considerations for implementing this technique in future studies.

However, at the <u>level of individual participants</u>, only 30% showed significant effects of SL in their neural response, and among them the effects did not occur consistently at the same frequencies/harmonics. Conversely, performance on the implicit target-detection task administered post-exposure demonstrates that SL occurred in a substantially larger proportion of individuals (70%). Hence, the current results suggest that the EEG-based metric has a lower sensitivity than some implicit behavioral metrics, and likely underestimates the prevalence of SL in individual participants.

### Strengths and weaknesses of the frequency-tagging approach for assessing SL

The EEG frequency tagging approach has been proposed as a more direct means for assessing SL, circumventing the need for behavioral post-exposure testing. Among its strengths is its online nature, that allows researchers to track the formation of a neural representation for pseudowords over time, without introducing a dual task. This approach has been successfully applied for studying neural processing of familiar and unfamiliar languages, and how the representation of different linguistic levels of speech is modulated by factors such as attention, state of arousal and consciousness (Ding et al., 2016; Makov et al., 2017; Getz et al., 2018; Niesen et al., 2019; Chen et al., 2020; Luo and Ding, 2020; Har-shai Yahav and Zion Golumbic, 2021). The frequency-tagging approach has also brought great excitement to the field of Statistical Learning, since it offers a way to dissociate between the acoustic-representation of individual elements in a stream (e.g. syllable rate, 4Hz in the current study) and its parsing into larger units (e.g. pseudoword rate; 1.33Hz and its harmonics in the current study) that reflects higher-level generalization and learning (Buiatti et al., 2009; Batterink and Paller, 2017, 2019; Getz et al., 2018; Elmer et al., 2021; Henin et al., 2021).

However, here it is important to note an important methodological caveat: The interpretation that peaks in the neural response at the pseudoword rate reflect detection and parsing of pseudowords relies on the assumption that these peaks cannot be derived from the acoustics of the stimulus alone. Unfortunately, this assumption does not seem to hold for the type of stimuli used in the triplet-based Artificial Language SL paradigm. As shown in the modulation spectra when testing several different combinations of triplet-syllables, a prominent peak can be seen at the triplet-rate in addition to the syllable rate. This peak is generated due to subtle yet systematic differences in the envelope-shape of different syllables, that are presented consistently at the same position – an inherent feature of pseudowords. These caveats of the frequency-tagging approach have recently been pointed out when using bi-syllabic words in real languages (Luo and Ding, 2020; Har-shai Yahav and Zion Golumbic, 2021). Similarly, for Artificial Languages, the elegant re-analysis of the data in Batterink & Paller (2017) showed that at least part of the neural response at the triplet-rate frequency can be attributed to differences in the Obligatory Contour Principle (OCP)-place of different syllables rather than SL per se (van der Wulp, 2021). Therefore, in order to avoid overinterpretation of these peaks, adequate controls must be implemented in all studies.

Here we addressed this concern by introducing a Position-Controlled Baseline stimulus, that shared the same modulation spectrum as the Artificial Language stimulus. As expected, in addition to the neural response at the syllable rate, the response to this Position-Controlled stimulus contained a prominent peak at the triplet-rate and its harmonics, even though it contained no statistical regularities. This demonstrates the methodological caveat of frequency-tagging mentioned above – that the mere existence of a triplet-rate peaks is not, in and of itself, indication for Statistical Learning. Nonetheless, when comparing the neural response to the two stimuli at the group level, the triplet-rate peak (and its harmonic) was significantly larger in response to the Artificial Language stream relative to its Position-Controlled Baseline stimulus. This pattern suggests that the neural response at the triplet-rate and its harmonics reflects *a combination* of acoustic responses as well as responses reflecting detection of the underlying statistical structure and/or pseudoword boundaries.

Interestingly, the strongest effect was not found at 1.33Hz, which is the triplet-rate itself, but rather at its 3^rd^ harmonic (5.33 Hz). This is similar to the pattern reported by a recent ECoG study, where the most prominent effects of SL were also found at harmonics of the triplet-rate (Henin et al., 2021). Moreover, as detailed below, when inspecting the individual-level spectra, we found great variability in which frequencies showed the most prominent SL effects. The manifestation of effects at harmonic-frequencies is a natural consequence of presenting rhythmic stimuli, and should not necessarily be interpreted as carrying nuanced information regarding the nature of neural encoding for these stimuli (Zhou et al., 2016). However, this variability does present another potential caveat for the utility of the frequency-tagging approach.

### Assessing SL in Individual participants

One of the main goals of the current study was to investigate the sensitivity of different measures of SL at the level of individual participants. Due to the proposed ubiquitous nature of SL and is proposed importance for language acquisition, we expected to find evidence for SL in most participants. However, this was not case. Rather, the pattern emerging from comparing the three independent measures used here – the explicit 2AFC, the implicit target-detection task, and the frequency-tagged EEG spectrum – illustrates the operational challenge of empirical assessment of SL. The 2AFC test failed to show a significant effect at the group level, and at the individual level only 3 participants (7.5%) showed significant effects. The poor performance-levels are in line with previous studies where reported group-level detection rates range between 54%-74%, and individual-level significance rates are low (fewer than 50% of participants) (Franco et al., 2015b). This task also has been shown to have a medium-low test-re-test reliability (Siegelman and Frost, 2015; Erickson et al., 2016), and several methodological factors have been proposed explaining the low sensitivity of the 2AFC approach (Siegelman et al., 2017). There also seems to be a lack of correlation between the various auditory SL tasks themselves. A study comparing several auditory SL paradigms using the explicit 2AFC task on the same participants reported a lack of correlations between these very similar paradigms that only differed in the language that was used (Erickson et al., 2016). The authors therefore concluded that these low correlations were most likely the result of the poor psychometric properties of the 2AFC measure and that using a composite score of all these measures combined gives the clearest picture of the situation. Given these low performance rates, which do not coincide with other measures, it seems that this metric is not sufficiently reliable for determining whether SL has or has not occurred in individual participants.

The weakness of explicit 2AFC testing has led to the development of more implicit measures for assessing Statistical Learning. Some examples of implicit tasks include the target detection task used by (Batterink et al., 2015; Batterink and Paller, 2017) and adapted in the current study, as well as the Rapid Serial Auditory Presentation (RASP) (Franco et al., 2015a), statistically induced chunking recall (SICR), (Isbilen et al., 2017, 2020), and the click detection task (Gómez et al., 2011; Franco et al., 2015b). These tasks rely on a similar principle; if pseudowords in the stream are learned, this will produce a faster implicit response to targets that are associated with that pseudoword. In the current study, the implicit target-detection test was by far the most sensitive measure of SL, with 19/27 participants (70%) showing a significant effect on *either* hit-rate or RT. However, only 3 participants had significant effects in *both* measures, perhaps reflecting speed-accuracy tradeoffs, thus diluting the group-level effect of both measures. Nonetheless, the fact that *some* effect was observed in a high percentage of participants supports the assertion that most of them did pick up on the underlying regularities of the Artificial Language. At the same time, it is important to note that this high-proportion of individual-level effects is not found in all studies. For example, Batterink et al. (2015) report that 43% of participants demonstrated behavioral effects reflecting SL, in a target detection task similar to the one used in the current study. Two studies using a similar click detection tasks found vastly different proportions of individual-level effects, with Gómez et al. (2011) reported effects consisted with SL in 85% of participants, Franco et al. (2015b) found these in only 35% of participants, and many actually showed reverse effects. Taken together, although in the current study the implicit target-detection task seemed to be in line with the proposed ubiquitous nature of SL, the large variability across behavioral studies (in methods and results) makes it difficult to wholeheartedly accept these implicit measures as a reliable benchmark for assessing SL. Moreover, the cross-study discrepancies make it extremely difficult to determine the true extent of SL in individual participants.

The diverse and inconclusive nature of indirect behavioral measures was one of the primary motivators for looking to neural measures as more direct signatures of SL. The current study is the first to assess the robustness of neural SL measures in individual participants using the frequency-tagging approach. In contrast to the expected ubiquity, we found that only 12/40 participants (30%) showed significant effects of SL in their EEG spectra. One reason for this might be the poor signal-to-noise ratio (SNR) in individual-level scalp level EEG. Indeed, a recent ECoG study, which by its nature is based on individual participants, was able to demonstrate robust neural response at pseudoword-related frequencies, suggesting that improving the signal SNR might lead to more robust results (Henin et al., 2021). However, another factor that exacerbates the complexity of interpreting the frequency-tagging results is that the frequencies where SL effects were observed varied substantially across participants, with effects found either at the pseudoword rate or the 1^st^ or 2^nd^ harmonic. This was the case also in the ECoG data reported by Henin et al. (2021), which leaves many questions open regarding the underlying mechanism. We can hope that future methodological advances will improve the SNR of frequency-tagging measures, which in turn might reveal more extensive evidence for SL. However, at present, the current results leave us wondering whether the low prevalence of neural-effects corresponding to SL are merely a result of poor SNR or if they challenge the assumption of the ubiquitous nature of SL.

In the absence of a ‘ground truth’ indication for SL, we turn to look for evidence of converging operations among the multitude of tests, that all supposedly measure whether SL has taken place, relate to each other. Unfortunately, results from the different behavioral and neural measures do not seem to converge as one might expect if they truly capture the same cognitive operation. In testing whether neural results corresponded in any way with the behavioral responses, we found that the subgroup of participants who showed neural evidence for SL also had slightly faster RTs in the implicit target detection task than those who did not. However, no correspondence was found when examining the with-participant correlation, nor were there any correlations with other behavioral measures. The current results align with previous studies that also reported no correlation between results on explicit and implicit methods of testing for SL (Misyak et al., 2010; Batterink et al., 2015; Franco et al., 2015a; Isbilen et al., 2020). In the few studies where there were significant correlations between explicit and implicit measures, these were not consistent across different modalities (Isbilen et al., 2020), or differences in the explicit task (Batterink and Paller, 2017). Some have opted to interpret the lack of a reliable cross-measures correlation as an indication that each measure picks up on a different cognitive-aspect of SL, e.g., suggesting a dissociation between explicit recall and implicit learning (Batterink et al., 2015; Franco et al., 2015a; Isbilen et al., 2017). However, we cannot rule out the possibility that all of these measures – behavioral and neural alike - are simply too crude or too indirect for assessing the formation of internal memory representations arising from SL. Consequently, it seems that we still lack a ‘ground truth’ indication for SL, which (at the moment) severely limits the extent to which this ability can be studied at the level of individual participants.

## Conclusions

The current study highlights the utility and the limitations of the EEG frequency-tagging approach as a research tool for studying SL. At the group level, our results seem promising, providing evidence that observed peaks in the neural signal at the pseudoword frequency (and its harmonics) are probably not only a result of acoustic artifacts, but also reflect detection of underlying transitional-probabilities. However, the frequency-tagging approach might not be as useful for studying SL in individual participants, given its low sensitivity, the fact that it does not always manifest at the same frequencies, and its overall low correspondence with behavioral tests of SL. Therefore, while some researchers may find this experimental approach suitable for their needs, these limitations and potential confounds should be taken into consideration when interpreting and comparing results across studies.

## Funding sources and Acknowledgements

This work was supported by the German-Israel Foundation, grant # 1422. We would like to thank Prof. Lucia Melloni for helpful conversations during the conceptualization and execution of this project.

